# Hippocampal sharp-wave ripples decrease during physical actions including consummatory behavior in immobile rodents

**DOI:** 10.1101/2024.12.26.630368

**Authors:** Tomomi Sakairi, Masanori Kawabata, Alain Rios, Yutaka Sakai, Yoshikazu Isomura

## Abstract

Hippocampal sharp-wave ripples (SWRs) are intermittent, fast synchronous oscillations that play a pivotal role in memory formation. It has been well-established that SWRs occur during “consummatory behaviors”, e.g., eating or drinking a reward for correct action. However, most of typical behavioral experiments using freely moving rodents have not rigorously distinguished between the act of eating/drinking (regardless of consummation or consumption) from stopping locomotion (immobility). Therefore, in this study, we investigated the occurrence of SWRs during a reward-seeking action and subsequent consummatory reward licking in constantly immobile rats maintained under head fixation and body covering. Immobile rats performed a pedal hold-release action that was rewarded with water every other time (false and true consummation). Unexpectedly, the SWRs remarkably decreased during reward licking as well as pedal release action. Unlearned rats also showed a similar SWR decrease during water licking. Conversely, SWRs gradually increased during the pedal hold period, which was enhanced by reward expectation. A cluster of hippocampal neurons responded to cue/pedal release and reward, as previously shown. Some other clusters exhibited spike activity changes similar to the SWR occurrence, i.e., decreasing during the pedal release action and reward licking, and enhanced by reward expectation during pedal hold period. These task event-responsive neurons and SWR-like neurons displayed stronger spiking synchrony with SWRs than task-unrelated neurons. These findings suggest that the hippocampus generates SWRs, which may associate action with outcome, in “relative immobility” (action pauses) rather than specific consummation or consumption.

**Significance Statement:** To clarify the characteristics of hippocampal sharp-wave ripples (SWRs), we analyzed the SWRs occurring during operant task performance in immobile rats under both head fixation and body covering. First, we found that SWRs decreased when they licked and drank water, conflicting with the theory that SWRs occur in consummatory behavior. Second, hippocampal neurons showed different task-related activities, particularly those that resembled SWR occurrences or conveyed specific signals on task events. Third, these task-related neurons displayed strong synchronous discharges during SWRs in task-engaged periods. These findings may explain the neuronal mechanisms underlying the association between an action and its outcome.

## Introduction

Human and animal behaviors can be psychologically explained as either *appetitive* (*preparatory*) behaviors, which seek appetitive, sexual, or sleep desire; or *consummatory* behaviors, which satisfy desire (Craig, 1918; Woodworth, 1918; Konorski, 1967; Buzsáki, 2015). For example, when we are thirsty, filling a glass of water from the faucet is a preparatory behavior, while drinking the water is a consummatory behavior (NB, this term originally meant “consummation” in psychology and ethology, but is also used to indicate “consumption”). The hippocampus, which is responsible for episodic memory formation, also plays a role in operant action-outcome learning in rodents (Corbit and Balleine, 2000), including preparatory and consummatory behaviors (Osborne and Dodek, 1986; Flaherty et al., 1998; Jurado-Parras et al., 2013). In fact, when rats operantly learn to release a pedal (preparatory behavior) to drink water as a reward (consummatory behavior), hippocampal neurons become responsive to the reward-seeking and reward-receiving phases (Soma et al., 2023).

In the hippocampus, high-frequency (100–250 Hz) synchronous oscillations, called “ripples,” occur intermittently, accompanied by a brief (40–100 ms) “sharp wave” in local field potentials (sharp-wave ripple, SWR; Buzsáki et al., 1983, 1992; Buzsáki, 1986). Hippocampal pyramidal cells and interneurons discharge synchronously at different phases of SWR oscillation (Ylinen et al., 1995; Klausberger et al., 2003). SWRs originate from the hippocampal CA3 area, spread bilaterally throughout the CA1 area (Csicsvari et al., 2000), and propagate into the entorhinal and neocortical areas (Chrobak and Buzsáki, 1996), where they are associated with cortical activity (Sirota et al., 2003; Isomura et al., 2006). It is well known that SWRs occur during consummatory behaviors, such as eating and drinking (Buzsáki et al., 1983; Buzsáki, 1986, 2015), in addition to immobility (standstill) and slow-wave sleep, which can be regarded as consummatory behaviors in the broad sense (Vanderwolf, 1969; Hartse et al., 1979; Suzuki and Smith, 1985; Buzsáki et al., 1983, 1992; Buzsáki, 1986; O’Neill et al., 2006; Vöröslakos et al., 2023: for review, see Buzsáki 2015; Buzsáki and Tingley, 2023). On the other hand, continuous theta (4–12 Hz) oscillations appear in place of SWRs during whole-body movements (e.g., running) and paradoxical sleep (Vanderwolf 1969; Buzsáki et al., 1983; Buzsáki and Moser, 2013). When animals freely explore a space, a set of hippocampal pyramidal cells is sequentially activated along their receptive places during theta oscillations (place cells). Immediately after animals stop running, the place cells are instantly reactivated in reverse order during SWRs (replay; Foster and Wilson, 2006; Diba and Buzsaki, 2007). Importantly, the occurrence of SWRs and replays is facilitated by the presence of a reward (food or water) at the end of the spatial exploration (Singer and Frank, 2009; Ambrose et al., 2016; Sasaki et al., 2018; Sosa and Giocomo, 2021). These observations support the idea that running for exploration is a preparatory behavior for seeking a reward, whereas stopping at the reward leads to a consummatory behavior during which SWRs occur in the hippocampus, depending on the availability of the reward (either expected or acquired). However, most behavioral experiments conducted in freely moving animals have failed to rigorously distinguish between immobility (stopping) and reward intake (eating/drinking), even though it is necessary to determine whether reward expectation or acquisition facilitates SWR occurrence. Therefore, in the present study, we investigated how reward expectation and acquisition separately influence SWR occurrence during preparatory and consummatory behaviors in immobile rats under head-fixed conditions. Head fixation constantly avoids whole-body movements and allows for more precise analyses of behavioral events (pedal release, licking, etc.) than freely moving conditions. We recorded SWRs and neuronal spike activity in the hippocampal CA1 area while rats performed an operant pedal release task (Soma et al., 2017, 2019, 2023) requiring alternating rewarded and unrewarded actions (Isomura et al., 2013; Yoshizawa et al., 2023). In contrast to the accepted theory, we found that SWRs and some neuronal activities decreased during the consummatory reward-intaking action in immobile rats.

## Materials and Methods

### Animals

Protocols for all experiments were approved by the Animal Research Ethics Committee of Tamagawa University (H28-32, H30-32) and the Animal Care and Use Committee of Tokyo Medical and Dental University (currently, Institute of Science Tokyo; A2019-274, A2021-041, A2023-116), and were performed in accordance with the Fundamental Guidelines for Proper Conduct of Animal Experiment and Related Activities in Academic Research Institutions (Ministry of Education, Culture, Sports, Science, and Technology of Japan (MEXT)) and the Guidelines for Animal Experimentation in Neuroscience (Japan Neuroscience Society). All surgical procedures were performed with appropriate anesthesia (isoflurane or other, detailed below), and all efforts were made to minimize suffering. The animal experimental procedures have been described in our previous studies (Isomura et al., 2013; Kimura et al., 2012; 2017; Saiki et al. 2018; Soma et al., 2017, 2019, 2023; Rios et al., 2019, 2023).

Adult Long-Evans rats (Institute for Animal Reproduction; *N* = 9, 159–312 g, male and female) were maintained in their home cages under an inverted light schedule (lights off at 9:00 a.m.; lights on at 9:00 p.m.). They were handled briefly (10 min, twice) in advance to adapt to the experimenter and experimental environment.

### Surgery

For head-plate implantation, rats were anesthetized with isoflurane gas (4.5% for induction and 2.0–2.5% for maintenance; Pfizer) using an inhalation anesthesia apparatus (Univentor 400 anesthesia unit, Univentor), and were placed on a stereotaxic frame (SR-10R-HT, Narishige). The skins around surgical incisions were coated in Lidocaine jelly (AstraZeneca) to induce local anesthesia. During anesthesia, the rectal temperature was maintained at 37 °C using an animal warmer (BWT-100, Bio Research Center). The head-plate (CFR-1, Narishige) was attached to the skull using small stainless-steel anchor screws (M1, 2 mm in length) and dental resin cement (Super-Bond C&B, Sun Medical; Panavia F2.0, Kuraray Medical; Unifast II, GC Corporation). The reference and ground electrodes (Teflon-coated silver wires, A-M Systems; 125 μm in diameter) were implanted on the dura matter above the cerebellum. The skull surface was covered with silicone sealant (DentSilicone-V, Shofu). Analgesics and antibiotics were administered postoperatively, as required (meloxicam, 1 mg/kg *s.c.*, Boehringer Ingelheim; gentamicin ointment, 0.1%, *ad us.ext.*, MSD).

Following full recovery from this primary surgery (≥5 days later), the rats were allowed *ad libitum* access to water during the weekends, but obtained water only when they performed the task correctly during the rest of the week. When necessary, agar blocks containing 15 ml water were provided to the rats in their home cages to maintain them at >80% of original body weight (Soma et al., 2017, 2019, 2023; Rios et al., 2019, 2023).

### Behavioral task

In preliminary experiments for natural drinking (see **Fig. 1**), two naïve (unlearned) rats were held immobile with the head fixed to a stereotaxic frame and the body covered with a cylinder-like stainless-steel shelter (6 cm in diameter) using the task control system for head-fixed rodents (TaskForcer; O’HARA & Co., Ltd). In this situation, rats were “immobile” in the sense that they remained in place and did not involve locomotion, but they were able to flexibly move their limbs, change their posture and quickly relax in the cylinder-like shelter (Isomura et al. 2009; Kimura et al., 2012). A spout was placed in front of their mouth to dispense a drop of sweet water (0.1% saccharine, 5 µl/drop) from a micropump. Saccharin is not metabolized by rats, and has a negligible effect on blood glucose levels. Therefore, we did not consider the interactions between SWRs and glucose metabolism (Tingley et al. 2021). Their licking movements with the tongue were monitored by detecting the strain of the spout (KFG-2N strain gauge and DPM-911B amplifier, Kyowa), which was confirmed with an infrared video camera. The rats passively received two drops of water (10 µl at 150 ms interval) that were delivered from the spout repeatedly every 10–20 s (0.1 s step at random; about 100 times in total) without any prior cues; hence, it was neither classical nor operant conditioning. The hippocampal activity was recorded electrophysiologically (see below) during the passive drinking performance of the two rats (*n* = 6 and 5 sessions).

**Figure 1.**
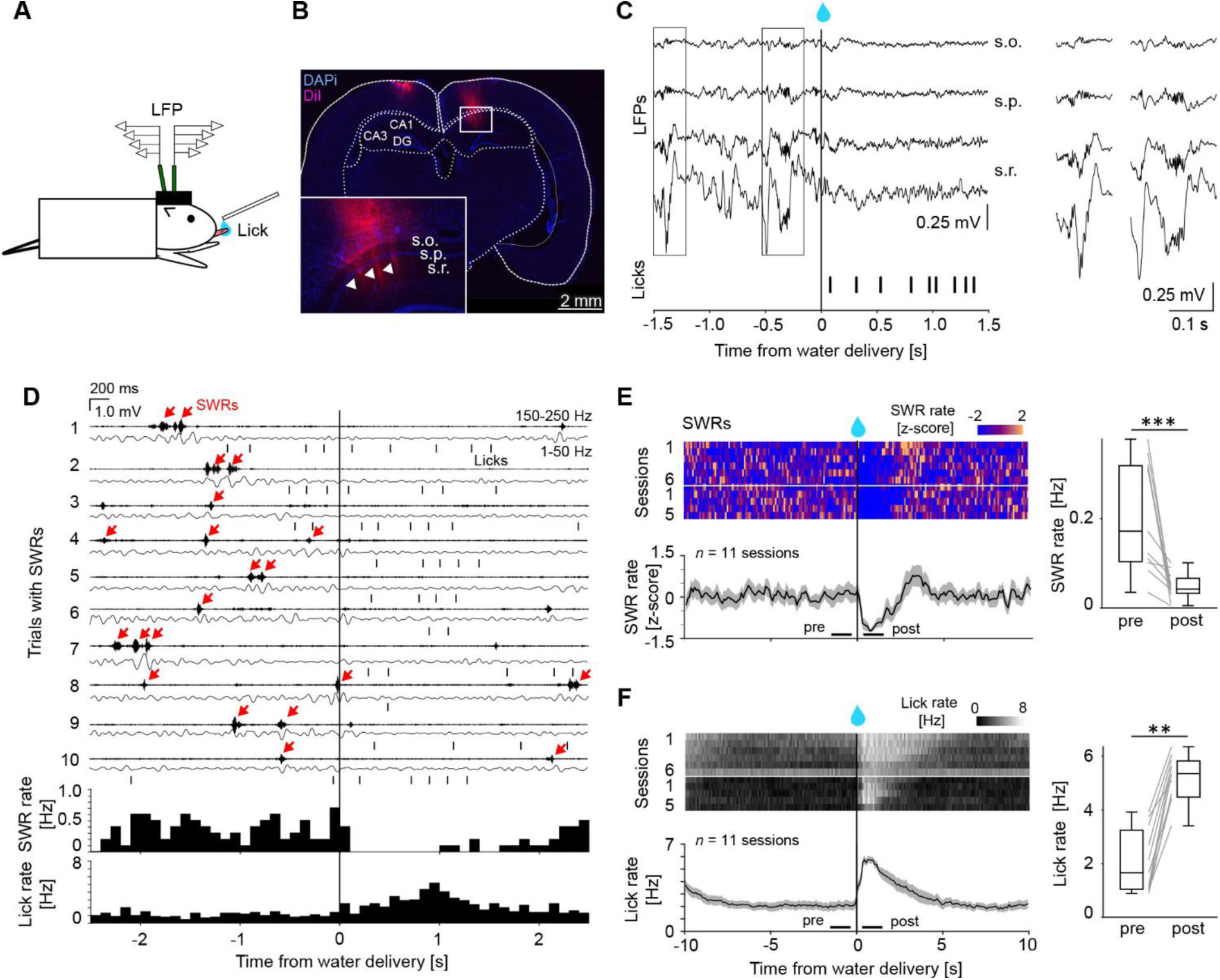
Hippocampal sharp-wave ripples (SWRs) disappeared during water drinking in naïve (unlearned) immobile rats. ***A***, Schematic diagram of electrophysiological recordings from the bilateral hippocampi of a head-fixed rat licking a drop of water from a spout (10–20 s, at random intervals). LFP, local field potential. ***B***, Silicon probe tracks in the hippocampal CA1 area (arrowheads, stained with DiI on the probe shanks). s.o., stratum oriens; s.p., s. pyramidale; s.r., s. radiatum. ***C***, *Left*, Two SWR events before water delivery (0 s) in raw LFP traces. *Right*, SWR traces enlarged from the left rectangles (ripple oscillations and sharp waves at s.p. and s.r., respectively). Note that there was no artifact noise caused by the licking (vertical symbols). ***D***, *Top*, Ten consecutive LFP traces with SWRs (red arrows) detected within ±2.5 s of water delivery (0 s), filtered at ripple (*each upper*, 150–250 Hz) and sharp wave (*lower*, 1–50 Hz) bands. Vertical symbols indicate the licks. *Bottom*, SWR (*upper*) and lick (*lower*) rates in this session (98 trials). ***E***, *Left*, Summary of z-scored SWR rate change (*upper*, individual sessions; *lower*, the mean and s.e.m.) around water delivery (0 s) in all sessions (11 sessions from 2 rats). *Right*, Box- and-whisker plots showing a decrease in SWR rate after water delivery (compared between pre and post 1 s windows). Asterisks represent statistical significance by Wilcoxon signed-rank test. \**p* < 0.05, \*\**p* < 0.01, \*\*\**p* < 0.001 in all figures. ***F***, *Left*, Summary of lick rates in all sessions. *Right*, Increase in lick rate after water delivery.

In main experiments on behavioral consummation (see **Fig. 2*A*,*B***), we have established “False-True Consummation (FTC)” task (controlled by the TaskForcer system), in which immobile rats operantly learned to hold and release a pedal with their forelimb (preparatory action) to obtain water as a reward by licking (consummatory action). This was a modified version of our pedal release task for head-fixed rats (Soma et al., 2017, 2019, 2023; Rios et al., 2019, 2023). In this task, the head-fixed rats (*N* = 7 rats; *n* = 59 sessions) spontaneously started each trial by holding down the left and right pedals with the corresponding forelimbs for 1–2 s (0.1 s step at random). The position of each pedal was continuously monitored using an angle encoder (hold position: lower 0–20% of full pedal range). After completing the pedal hold time, rats had to release the left pedal quickly (within 0.9 s) in response to the presentation of the Go cue (10 kHz pure tone for 0.1 s) for the reward, while keeping the right pedal held (0.9 s). The reaction time was defined as the time from the onset of the Go cue until pedal release just started within the hold position.

**Figure 2.**
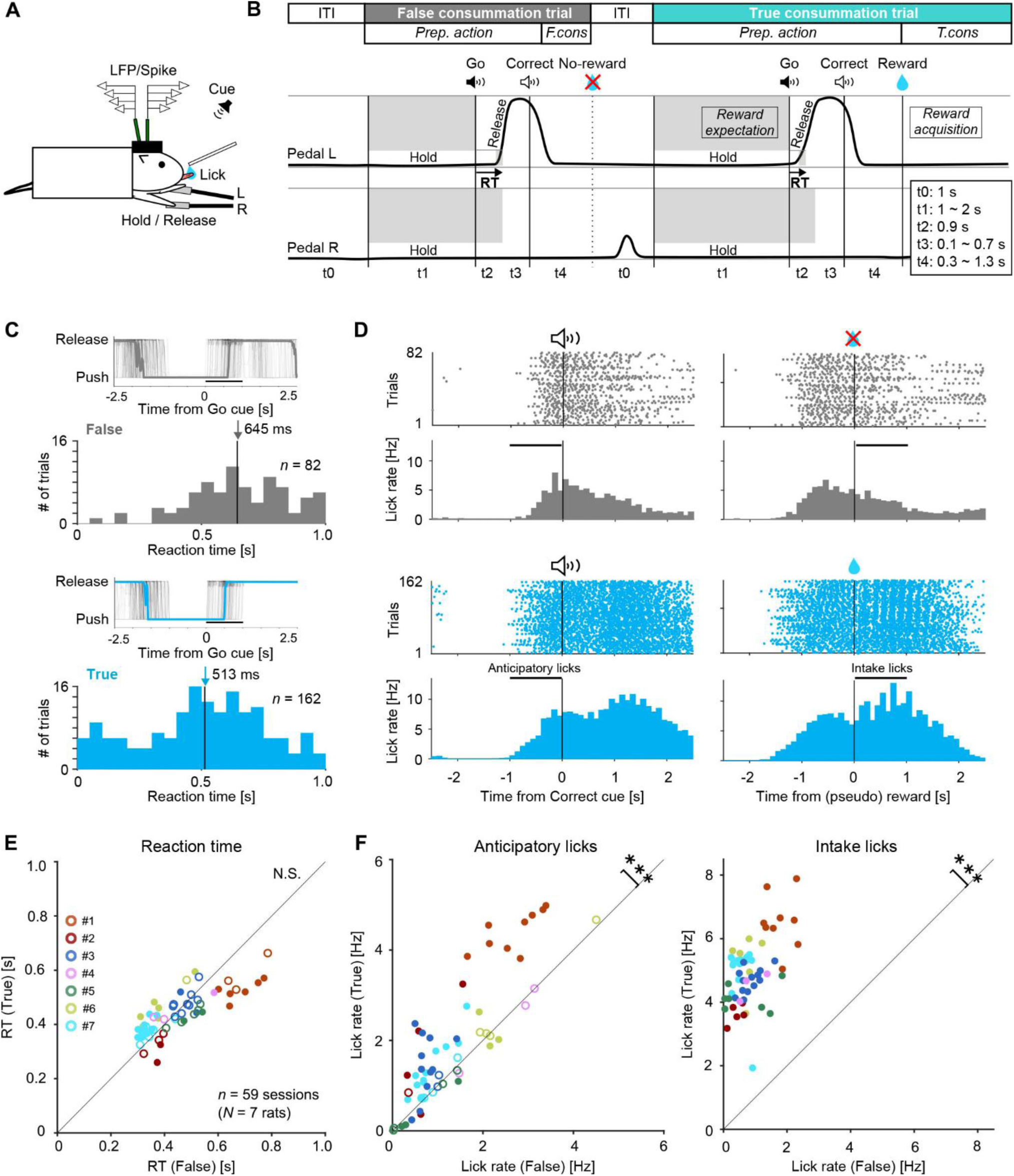
Behavioral performance of the False-True Consummation (FTC) task by immobile rats. ***A***, Pedal hold-release action in response to the auditory Go cue for reward water. ***B***, The FTC task comprised repetitive pairs of false (unrewarded) and true (rewarded) consummation trials. ITI, intertrial interval; Prep., preparatory action; F./T. cons., false/true consummatory action; RT, reaction time. The gray shaded areas indicate the forbidden range of the pedals. Nothing happened for no reward delivery after the Correct cue. See **Materials and Methods**. ***C***, Pedal traces (*upper*, all and median traces) and distribution of reaction times (*lower*; arrow indicates median) for false (*gray*) and true (*blue*) consummation trials in a session. We analyzed only correct trials without errors following previous trials. ***D***, Lick events in individual trials (*upper*) and lick rates (*lower*) aligned with the onset of the Correct cue (*left*) and no (pseudo)-reward/reward delivery (*right*). The horizontal bars indicate the time window for anticipatory and intake lick counts. ***E***, Comparison of the average reaction time between false and true consummation trials in all sessions (59 sessions from seven rats). Colored circles represent individual rats (filled shows statistical significance for each session). Group analysis showed no significant difference. ***F***, Comparison of the average lick rates between false and true consummation trials in all sessions. Anticipatory (*left*) and intake (*right*) lick rates were significantly higher in the true consummation trials.

If the rats were judged to react correctly, a Correct cue (8 kHz pure tone for 0.3 s) was presented to them 0.1–0.7 s (0.1 s step) after the correct judgement, and two drops of reward water (0.1% saccharine, 10 µl) were ready to be delivered from the spout another 0.3–1.3 s (0.1 s step) after the Correct cue presentation. However, the actual reward was given in every other trial, that is, alternating between reward and no-reward trials (reward alternation: Isomura et al., 2013; Yoshizawa et al., 2023). They could confirm that the action was correct by Correct cue presentation, regardless of whether it was rewarded or not. Such reward alternation allows rats to easily and naturally predict the reward or non-reward for the correct action (Tyler et al., 1953). In the present study, we considered a pair of no-reward and subsequent reward trials as “false consummation” and “true consummation” trials, respectively, for comparison (as described in **Results** and **Fig. 2*B***). If the rats moved the pedals incorrectly, an Error cue (3 kHz square-wave tone for 0.3 s) was presented to them immediately, and they were required to retry the same false or true consummation trial (correction trial). If they failed to react to the Go cue (omission), they also retried the correction trial. We analyzed only correct test trials without errors following the previous trials to ensure their reward expectation. They proceeded to the next trial in a self-paced manner after an intertrial interval (ITI; 1 s). The rats typically learned this operant task in one month (2–3 h training a day), reaching the analysis criteria with ≥30 correct trials a session (42–566 trials/session).

### Electrophysiology

Once the rats recovered from the primary surgery (two rats in the preliminary experiments) or completed task learning (seven rats for the main experiments), they underwent a secondary surgery under isoflurane anesthesia for later electrophysiological recordings. We made two tiny holes (1.0–1.5 mm in diameter) in the skull and dura matter above the dorsal hippocampi bilaterally (3.8 mm posterior and ±2.0 mm lateral from the bregma). The skull surface and holes were immediately covered with antibiotic ointment and silicon sealant until the recording experiments.

We performed extracellular multichannel recordings of local field potentials (LFPs) and spike activity of individual neurons in the hippocampal CA1 area while the rats behaved under head fixation (11 preliminary sessions from two rats; 59 main sessions from seven rats). Two sets of custom-ordered 64-channel silicon probes (ISO64_4x_tet_lin_A64, NeuroNexus Technologies; 15 tetrodes and 4 tandem electrodes at 150 µm interval on 4 shanks with 50 µm thick) were precisely inserted into the hippocampal CA1 area bilaterally via a supportive layer of agarose gel (2% agarose-HGT, Nacalai Tesque) on the brain surface using fine micromanipulators (SMM-200B, Narishige). The two probes were angled 6°–9° towards the medial side to avoid mutual interference. The recording depth was determined so as to detect the ripple oscillation of the SWRs at the second tetrode from the tip of each probe, which corresponded to the stratum pyramidale (s.p.) layer of the hippocampal CA1 area (Csicsvari et al., 2000; Isomura et al., 2006). Hence, the upper and lower (tip) tetrodes were located in strata oriens (s.o.) and radiatum (s.r.), respectively. Probes were inserted at least 1 h prior to the start of each recording session. In the last recording session, the probes were coated with the red fluorescent dye DiI (DiIC18(3), PromoKine) to visualize the tracks later.

The multichannel signals were amplified by two sets of preamplifiers (MPA32I, Multi Channel Systems) and main amplifiers (FA64I, Multi Channel Systems; final gain 2,000, band-pass filter 0.5 Hz–10 kHz), and then digitized and stored at 20 kHz with a 128-channel recorder system (USB-ME128, Multi Channel Systems). The two pedal traces, licking (strain) signals, and task event signals (pedal hold and release, Go cue, Correct cue, reward, error, ITI, etc.) were digitized and stored simultaneously using the same system.

### Histology

After the final recording experiments, rats were deeply anesthetized with urethane (2–3 g/kg weight, *i.p.*), and transcardially perfused with cold saline followed by 4% paraformaldehyde in 0.1M phosphate buffer. The entire brain was postfixed, immersed in 30% sucrose in PB for cryoprotection, and sectioned coronally at 20 µm thick using a cryostat (CryoStar NX50, Epredia). We checked the red fluorescence of the probe tracks with DiI and hippocampal CA1 layers counterstained with DAPI (DAPI-Fluoromount-G, Southern Biotech) under a motorized fluorescence microscope (IX83-DP80, Olympus).

### SWR detection

SWR events were detected using MATLAB (MathWorks), primarily following the standard threshold criteria (Csicsvari et al., 1999, 2000; Isomura et al., 2006; O’Neill et al., 2006). In brief, the multichannel signals were down-sampled to 1 kHz and filtered at the ripple band (150–250 Hz) *offline*. The filtered signal was squared, smoothed with a Gaussian kernel (σ = 4 ms), square rooted, and z-scored through the entire session for each channel. The SWR candidates were detected by a threshold of four standard deviations (SD) from the baseline, and the duration was defined as > 4 SD. The SWRs shorter than 30 ms and longer than 200 m were excluded from the analysis. The peak amplitude and peak time of each SWR event were defined as the negative maximum and its time position of the ripple band LFP during the event. If the SWRs overlapped among the channels, the SWR with the largest amplitude was selected for analysis. Possible artifacts, if contaminated, were eliminated by visual and tailored inspection according to the artifact form, independent of the task information.

### Spike isolation and neuron classification

Multichannel data were also processed to isolate the spike events of individual neurons in each tetrode of the silicon probes. Spike candidates were detected and clustered using the spike-sorting software, EToS (Takekawa et al., 2010, 2012). Spike clusters were subsequently divided, combined, and discarded manually using the cluster curation software Klusters and NeuroScope (Hazan et al., 2006) to refine single-neuron clusters based on two criteria: the presence of refractory periods (>2 ms; spike contamination with less than 30% of ongoing spike rate) in their own autocorrelograms, and the absence of refractory periods in cross-correlograms with other clusters. We included single-neuron clusters if they exhibited a sufficient number of spike trains during task performance (≥200 trials with total ≥1,000 spikes).

For each neuron (spike cluster), we analyzed the basic spike properties and functional spike activity related to behavioral task performance using MATLAB software. Based on the bimodal distribution of spike duration (from trough to peak; Soma et al., 2023), individual neurons were classified as either regular-spiking (RS) (≥0.55 ms; mostly putative pyramidal cells) or fast-spiking (FS) (≤0.5 ms; putative inhibitory interneurons) (see **Fig. 4*A***) neurons. The ongoing spike rate was defined as the mean spike rate over the entire period when the neuron was recorded stably throughout the session.

For task-related activities, we analyzed spike trains in relation to two task events (Go cue and reward delivery) separately in the correct trial data. The spike trains were aligned with the timing (0 s) of these task events and analyzed in 2.0 s time window (−1.0 to +1.0 s from Go cue and −1.0 to +1.0 s from reward delivery). A no-reward event (denoted as “pseudo-reward” in the figures) was assumed to occur at the same time as the reward delivery event in each pair of false-true consummation trials for fair comparison in analysis. The task-related activity was defined by the two task relevant indices (TRI*d* and TRI*p*) using *d* and *p* values of Kolmogorov–Smirnov (KS) test (see **Fig. 4*B***; “task-related” if TRI*d* ≥ 0.1 AND TRI*p* ≤ 10^−5^ for ≥100 available spikes; see also Kimura et al., 2017; Saiki et al., 2018; Soma et al., 2017, 2019, 2023) for four combinations of the two task events [Go cue and reward/no-reward] with the two trial types [false and true consummation trials]. If a neuron was task-related in any of the four activity patterns, it was referred to as a task-related neuron; otherwise, it was referred to as a non-task-related neuron.

Task-related neurons were further classified into several neuron clusters (Clu.1–4) using the following spectral clustering procedure. For each neuron, spike rate patterns were calculated for −1 to +1 s from the Go cue and for −1 to +3 s from the reward delivery (10 ms bin) in true and false consummation trials. The four patterns were concatenated and z-scored as one 12 s trace. The correlation coefficients among all neurons were calculated to create a correlation coefficient matrix. Spectral clustering was performed using the correlation coefficient matrix as the similarity matrix (spectralcluster function in MATLAB; *k* = 4). The optimal number of clusters was selected as the smallest number of clusters for which the value of the smallest eigenvalue of the Laplacian matrix was stationary.

### Functional analysis of neural activity

We applied Phase-Scaling (PS) analysis (Kawabata et al., 2020) to determine which of the two task events (Go cue, pedal release, Correct cue, reward delivery, etc.) with varying inter-event intervals was temporally more relevant for neural activity such as the SWRs and neuronal spike activity. This analysis yielded the Phase index along the trial time normalized by the inter-event interval. Phase values closer to 0 and 1 indicate greater relevancies to the first event and second event, respectively. We can compute the Phase separately (P*−* for onset, P+ for offset) for the greatest increasing and decreasing trends in the activity change.

We also evaluated the differential changes in SWR occurrence and neuronal activity between false and true consummation trials using the reward modulation index (RMI), defined by the following equation (Isomura et al., 2013; Yoshizawa et al., 2023):

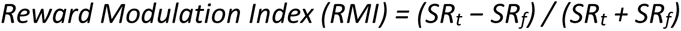

where *SR_t_* and *SR_f_* are the mean spike rates in the true and false consummation trials, respectively, in the time window before the Go cue (pre Go cue; 1 s), or after the no-reward/reward delivery (post reward; 2 s). If the RMI is >0, the activity is considered higher in true consummation trials than in false trials, and *vice versa*. The former RMI (pre Go cue) reflects modulation by reward expectation, while the latter RMI (post reward) reflects modulation by reward acquisition.

To evaluate the synchrony between the SWRs and spike activity of individual neurons, we made a cross-correlogram (CCG) of the spike sequence of each neuron with the SWR peak at time 0 s. For CCG analysis, the SWR events without other SWRs nearby (±0.5 s) were separately collected from the task-engaged (on-task) and ITI/resting (off-task) periods. We further analyzed the on-task SWRs in false and true consummation trials together or separately, but no further attempt was made to subdivide the on-task period into trial phases (e.g., pedal hold) because of the insufficiency of SWRs available for analysis. Spike increases or decreases during the SWRs were first discriminated by comparing the mean spike rate during the SWR time window (±50 ms from the SWR peak) with the overall (±0.5 s), and then judged statistically as a significant change or not by KS test (*d* ≥ 0.01 and *p* ≤ 10^−5^). The peak index of the spike rate during SWRs was defined as the difference between the two spike rates (true and false; on-task and off-task). The SWR-spike similarity was defined as the correlation coefficient (*r*) between the SWR rate and spike rate changes aligned with the task event (Go cue or reward).

### Statistics

Data in the text and figures are expressed as the mean ± SD or median with quartiles (unless otherwise mentioned) and sample number (*n*). We used appropriate statistical analyses: Wilcoxon signed-rank test, Wilcoxon rank sum test, Kruskal–Wallis test with post hoc Dunn test, KS test, and chi-square (*χ^2^*) test. The alpha value was set to 0.05 for statistical comparisons, except for using *p* value as a practical threshold to detect an activity change in the peri-event time histograms (PETH) and CCG data.

## Results

### Hippocampal SWRs during water licking in unlearned immobile rats

It has been well established that SWRs appear in the hippocampus of freely moving rodents during eating and drinking (Buzsáki et al., 1983; Buzsáki, 1986; Buzsáki and Tingley 2023). First, we conducted a preliminary experiment to check whether SWRs would appear when naïve (unlearned) rats drank water under an immobile condition of head fixation and lower body covering (**Fig. 1*A***). These immobile rats had never experienced any learning task before, and naturally licked and drank a drop of water with their tongues delivered from a spout approximately every 10–20 seconds. We recorded the SWR activity in the hippocampal CA1 area while monitoring licking behavior around the delivery of water. Two multichannel silicon probes (64-channels on four shanks) were placed bilaterally along the stratum oriens (s.o.), stratum pyramidale (s.p.), and stratum radiatum (s.r.) of the dorsal hippocampal CA1 area (**Fig. 1*B***). We clearly observed ripple waves (at the putative s.p. layer) and negative sharp waves (at the putative s.r. layer) simultaneously in the unfiltered local field potentials (**Fig. 1*C***). We also detected individual lick events using a strain-gauge sensor attached to the spout. The SWR and lick events were aligned with the timing of water delivery throughout all trials in this session (**Fig. 1*D***). The licking increased with a slight delay in water delivery. In contrast to our expectations, the occurrence of the SWR vanished during licking for water and gradually returned to baseline. We did not observe any prominent theta oscillations during licking (**Fig. 1*C***). Group data analysis also revealed that the SWRs abruptly disappeared in response to the water delivery from the first session, while the licking increased to drink the water (**Fig. 1*E***,***F***; *N* = 2 rats, *n* = 11 sessions: SWRs, pre 0.238 ± 0.118 Hz, post 0.076 ± 0.056 Hz, Wilcoxon signed-rank test, *p* = 1.28 × 10^−4^, *z* = 3.83, *r* = 0.578; Licks, pre 2.07 ± 1.18 Hz, post 5.17 ± 0.88 Hz, *p* = 3.34 × 10^−3^, *z* = −2.93, *r* = −0.626). Similar results were obtained under different conditions, where the water was delivered quickly every few seconds or where the rats had already sufficiently experienced a learning task (data not shown).

Consistent with our preliminary data, it has been reported that SWRs were reduced during reward licking in head-fixed mice performing a classical conditioning task (see Klee et al., 2021, its Figure Supplement 2B for Figure 5). These observations suggest that immobile animals may not necessarily display SWR generation with the same behavioral characteristics as those established in numerous studies on freely moving animals.

### Preparatory and consummatory behaviors in immobile rats

Figure 1 only shows that the SWRs decreased when water was passively administered to immobile rats. This raised the question as to whether such a decrease in SWRs also occurs in immobile rats while licking reward water as a consummatory behavior following explicit preparatory behavior. Another question is how the expectation of a reward affects SWR occurrence during preparatory behavior seeking the reward.

To answer these two questions, we trained head-fixed rats to perform an operant learning task (Soma et al., 2017, 2019, 2023), during which they obtained water as a reward (consummatory action) after correctly operating pedals with their forelimbs (preparatory action) (**Fig. 2*A***). The immobile rats started each trial by holding the right and left pedals with both forelimbs for 1–2 seconds (**Fig. 2*B***; t1). If they correctly released the left pedal quickly (within 0.9 s, t2) and kept holding the other pedal (for 0.9 s) in response to the presentation of the Go cue tone (0.1 s at 10 kHz), it was judged to be a correct trial. A Correct cue tone (0.3 s at 8 kHz) was presented to them 0.1–0.7 s (t3) after the correct judgement, and reward water was ready to be delivered from the spout another 0.3–1.3 s (t4) after the Correct cue presentation. Here, the actual reward was given every other trial, i.e., alternating between reward and no-reward trials (reward alternation: Isomura et al., 2013; Yoshizawa et al., 2023), allowing the rats to easily and naturally predict rewarded or unrewarded for the action (Tyler et al., 1953). In other words, rats must successfully complete a pair of no-reward and reward trials to obtain one reward (Fig. 2*B*; hereafter, “false” and “true” consummation trials, respectively). According to the psychological definition, it is a (true) consummation to lick and drink the reward water in a true consummation/reward trial. In contrast, attempts to lick the spout in the false consummation/no-reward trial are considered as false consummation, the only difference being the presence or absence of an actual reward. On the other hand, it is a preparatory action, in definition, to hold and release the pedals to seek the reward in both trials. The external task environment was identical between them; the only difference was the degree of reward expectation that the animals made internally. While the head-fixed rats were performing this “False-True Consummation” (FTC) task, we recorded the SWRs and spike activity in the hippocampal CA1 area bilaterally and monitored their pedal releases by each forelimb and their licks by the tongue. By comparing the false and true consummation trials, we examined how reward expectation and acquisition affected SWR generation during preparatory and consummatory actions, respectively.

The head-fixed rats usually learned to perform the FTC task in one month, and they completed several hundred pairs of false and true consummation trials in one session. In a typical session, the reaction time to release the pedal from the Go cue presentation was significantly shorter in true consummation trials than in false trials (**Fig. 2*C***), suggesting that the rats internally predicted the presence or absence of an upcoming reward when they released the pedal in each trial. Overall, the reaction time was not significantly different between the true and false consummation trials due to large individual differences (**Fig. 2*E***; *N* = 7 rats; *n* = 59 sessions: False 0.455 ± 0.127 s, True 0.444 ± 0.079 s, Wilcoxon signed-rank test, *p* = 0.624, *z* = 0.491, *r* = 0.064). The rate of “anticipatory licking” prior to the Correct cue presentation was significantly higher in true consummation trials than in false trials in most of the 59 sessions (**Fig. 2*D,F***, *left*; False 1.36 ± 1.00 Hz, True 1.93 ± 1.37 Hz, *p* = 1.26 × 10^−6^, *z* = −4.85, *r* = −0.631). As expected, the rate of “intake licking” to drink the actual reward from the spout in true consummation trials was remarkably higher than that in false trials (**Fig. 2D,*F***, *right*; False 0.84 ± 0.57 Hz, True 4.95 ± 1.04 Hz, *p* = 2.39 × 10^−11^, *z* = −6.68, *r* = −0.870). These results indicate that reward expectation was reliably reflected in the reaction time (in some sessions) and anticipatory liking rate.

### Hippocampal SWRs during preparatory and consummatory actions

We detected SWRs in the hippocampal CA1 area of immobile rats that performed the FTC task under the head-fixed condition. In a typical session, SWRs occurred during the pedal hold period but decreased rapidly after the Go cue presentation and pedal release (**Fig. 3*A***). SWR occurrence was much larger in the true consummation trials than in the false consummation trials during the pedal hold period. Moreover, SWR occurrence rapidly ceased after the Correct cue and the delivery of the real reward only in the true consummation trials (**Fig. 3*B***). Correct cues did not affect the false consummation trials. Group data analysis showed similar SWR increase during the pedal hold period, which was significantly enhanced in the true consummation trials (**Fig. 3*C***, *left*; *n* = 59 sessions: False-pre 0.263 ± 0.130 Hz and -post 0.172 ± 0.126 Hz, True-pre 0.433 ± 0.150 Hz and -post 0.230 ± 0.146 Hz: Kruskal–Wallis test *p* = 8.90 × 10^−17^, *χ^2^* = 77.8, *η²* = 0.242; post hoc Dunn test, False-pre vs. False-post *p* = 8.52 × 10^−4^, True-pre vs. True-post *p* = 1.42 × 10^−10^, False-pre vs. True-pre *p* = 3.12 × 10^−7^, False-post vs. True-post *p* = 0.0416, False-pre vs. True-post *p* = 0.194, False-post vs. True-pre *p* < 1.00 × 10^−100^). Abrupt disappearance of SWRs after reward delivery was also observed (**Fig. 3*C***, *right*; False-pre 0.216 ± 0.148 Hz and -post 0.286 ± 0.168 Hz, True-pre 0.245 ± 0.137 Hz and -post 0.0281 ± 0.0324 Hz: *p* = 2.16 × 10^−24^, *χ^2^* = 113, *η²* = 0.319; False-pre vs. False-post *p* = 0.0307, True-pre vs. True-post *p* < 1.00 × 10^−100^, False-pre vs. True-pre *p* = 0.171, False-post vs. True-post *p* < 1.00 × 10^−100^, False-pre vs. True-post *p* = 2.32 × 10^−13^, False-post vs. True-pre *p* = 1).

**Figure 3.**
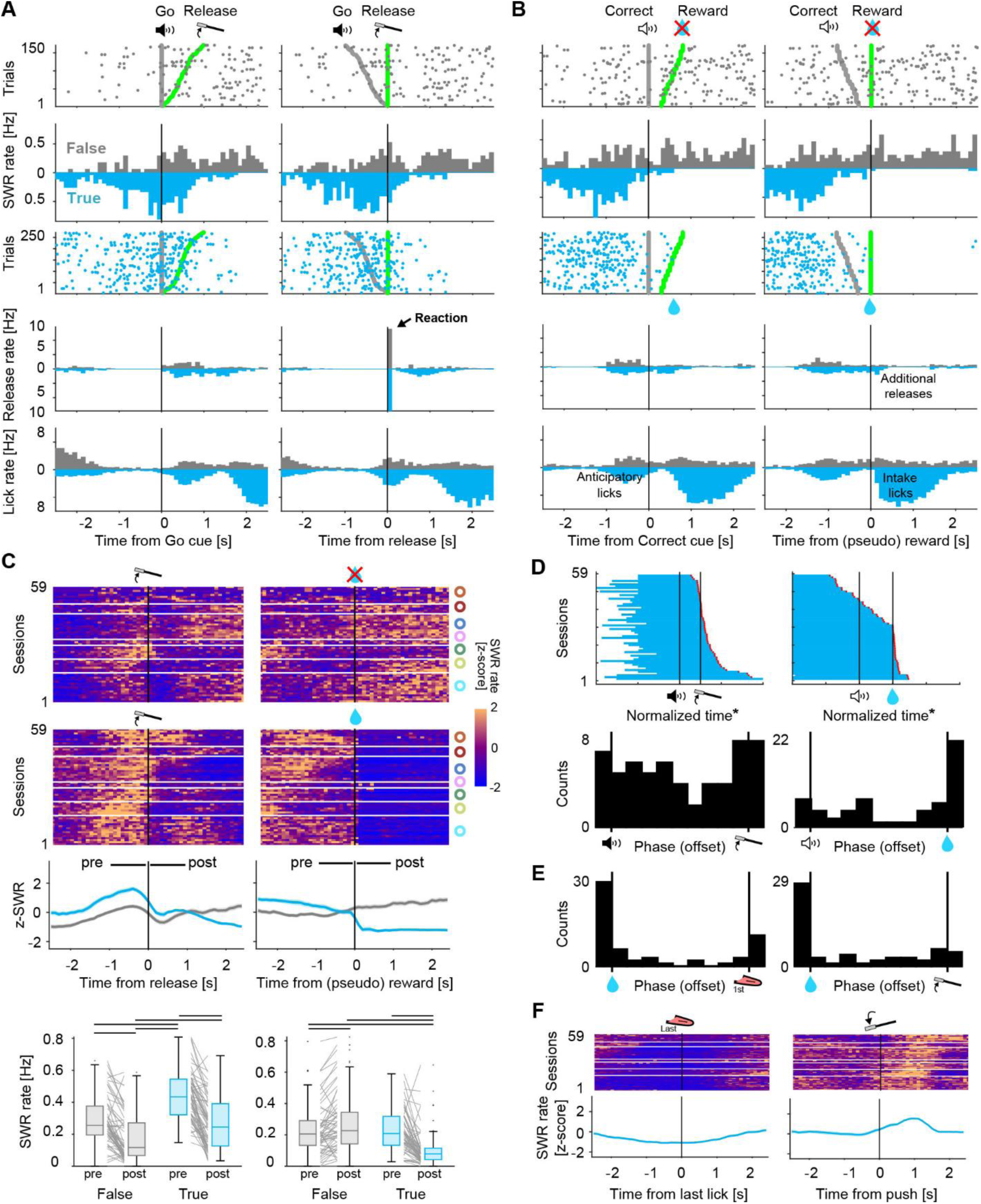
SWR occurrence during FTC task performance. ***A***, *Top*, SWR events in the individual trials (*upper* and *lower*) and SWR rates (*middle*) in the false (*gray*) and true (*blue*) consummation trials of a session, aligned with the Go cue (*left*; *thick gray*) and pedal release (*right*; *green*), and sorted by interval. *Bottom*, *upper*, Pedal release rate, including the correct reaction and additional pedal releases; *lower*, Lick rate. ***B***, SWR, pedal release, and lick rates aligned with the Correct cue (*left*; *thick gray*) and no (pseudo) reward/reward delivery (*right*; *green*). ***C***, *Top*, Summary of the z-scored SWR rate changes (*upper*, individual sessions for false and true consummation trials; *lower*, the mean and s.e.m.), aligned with the correct pedal release (*left*) and no-reward/reward delivery (*right*) in all 59 sessions. The colored circles indicate the rats in **Fig. 2*E***. Horizontal bars indicate the pre- and post-event time windows (1 s) for SWR counts before and after release (*left*), and no reward/reward (*right*). *Bottom*, Comparison of SWR rates among the pre and post events in false and true consummation trials. Horizontal lines indicate statistical significance (Kruskal–Wallis test with post hoc Dunn test). ***D***, Task events related to the decrease (offset) in the SWR rate, as determined by the Phase-Scaling (PS) analysis (Kawabata et al., 2020). *Top*, Onset-to-offset time-course of the SWR occurrence, temporally normalized (*) by the inter-event interval between the Go cue and release (*left*) and between the Correct cue and reward (*right*). Red edge indicates the offset. Note, however, that the SWR offset detection included the subsequent offsets (*left*, after release) and the preceding offsets (*right*, before reward), reflecting the two-step nature of the SWR decreases. *Bottom*, Phase distribution for the offset in the PS analysis. The Phase indicates the phase most temporally related to the activity offset. Note that the Phase for offset (i.e., SWR falling) was located between the Go cue and release (*left*) and immediately after the reward (*right*). ***E***, Task events related to SWR offset around the reward, showing a greater relation to the reward delivery itself than to the first lick (*left*) and first pedal release (*right*) following reward delivery. ***F***, Z-scored SWR rate (true consummation trials in all sessions) aligned with the last lick for reward intake (*left*) and the start of the pedal holding for the next trial (*right*).

Furthermore, we determined which of the two task events (Go cue or pedal release in **Fig. 3*A***; Correct cue or reward delivery in **Fig. 3*B***) was temporally more closely related to the SWR decrease in the true consummation trials using Phase-Scaling (PS) analysis (Kawabata et al., 2020). Regarding whether the Go cue or release, the Phase index for offset was bimodally distributed to 0 and 1 (**Fig. 3*D***, *left*), meaning that the SWR decreases were related to either of the two events, depending on the individual and the session. Regarding whether the Correct cue or reward, the Phase for offset was mostly 1 (**Fig. 3*D***, *right*), meaning that the SWRs disappeared at the time of the reward delivery rather than the Correct cue. Around the time of reward delivery, however, the rats moved their tongues for licking and might have additionally released the pedals to touch the spout or groom the face. Therefore, we further determined which of the reward delivery and the subsequent first lick and pedal release was most related to SWR disappearance (**Fig. 3*E***). This analysis clearly indicated that reward delivery (Phase 0 each) was the most plausible determinant of SWR disappearance around consummatory behavior. The SWR disappearance continued for one second after the end of the licking bout (**Fig. 3*E***, *left*), and then the SWRs gradually recovered towards the next pedal hold-release action (**Fig. 3*E***, *right*). As such, SWRs may begin to occur after the forelimb or tongue movement has completely stopped. Taken together, SWRs decreased during reward acquisition, that is, the consummatory licking action, in head-fixed rats.

### Hippocampal neuronal activity during preparatory and consummatory actions

We isolated spike activity from hippocampal CA1 neurons (*n* = 6,216) that were recorded simultaneously with SWRs during task performance. Based on the spike duration, they were classified into RS and FS neuron subtypes (**Fig. 4*A***; RS *n* = 4,794, FS 1,422). As predicted, the spike rate of RS (mainly, putative excitatory) neurons was statistically lower than that of FS (putative inhibitory) neurons (RS 1.57 ± 2.56 Hz, FS 3.78 ± 6.92 Hz; Wilcoxon rank sum test *p* = 3.53 × 10^−23^, *z* = −9.92, *r* = −0.10). We obtained task-related neurons using the TRI*d* and TRI*p* (**Fig. 4*B***; *n* = 3,213; see **Materials and Methods**). These task-related neurons were further classified into four neuron clusters (Clu.1 to 4) using spectral clustering for activity patterns in relation to the FTC task events (**Fig. 4*C***; Clu.1 RS *n* = 615, FS 85; Clu.2 RS 873, FS 124; Clu.3 RS 691, FS 163; Clu.4 RS 421, FS 241; see **Materials and Methods**).

**Figure 4.**
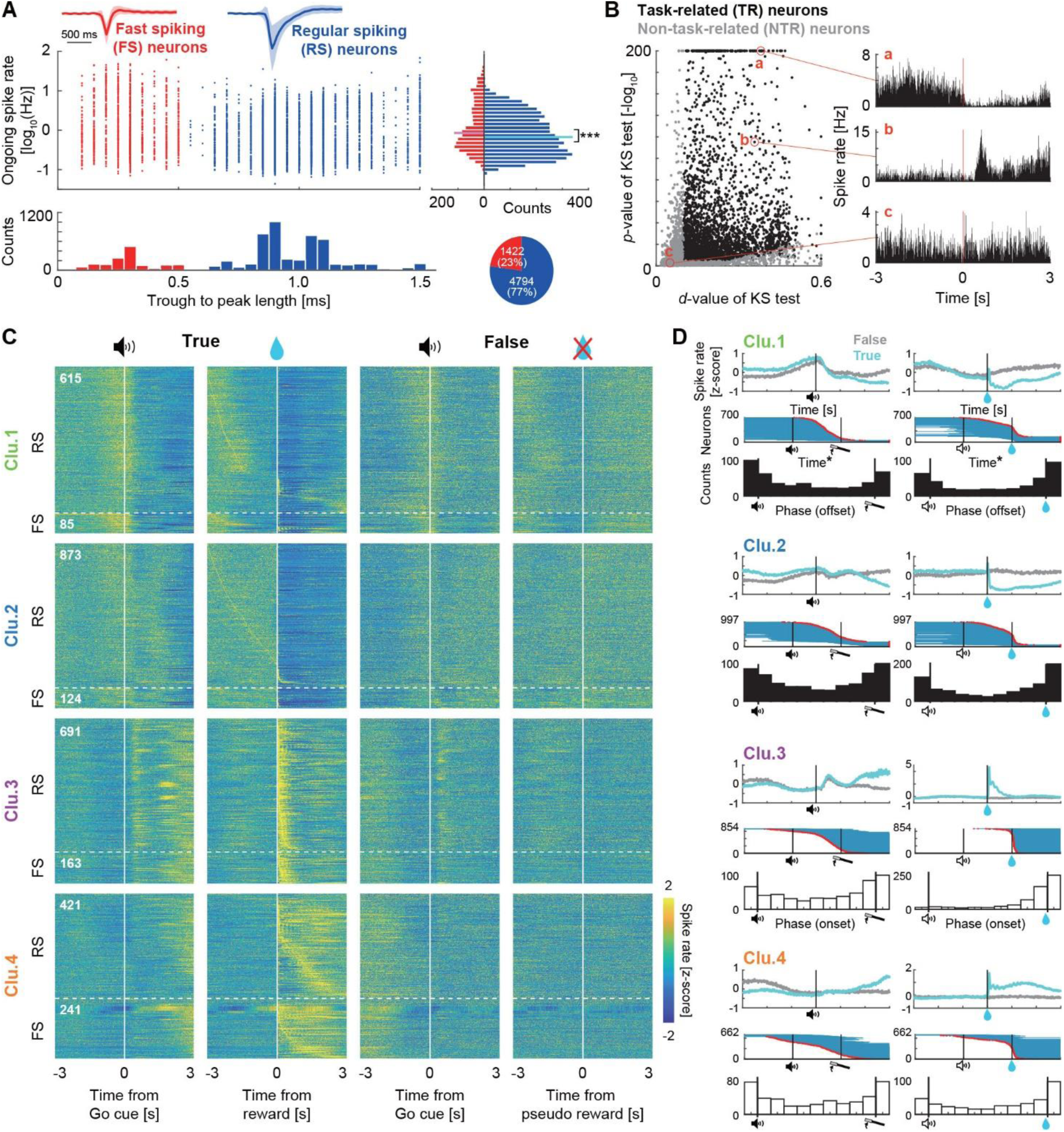
Spike activity of hippocampal neurons during FTC task performance. ***A***, Classification of all isolated neurons into regular spiking (RS; *blue*) and fast spiking (FS; *red*) neurons based on the spike waveform (trough-to-peak length). The ongoing spike rate (on a logarithmic scale; averaged over the entire recording time) was significantly lower in RS neurons than in FS neurons. ***B***, *Left*, Criteria defining the task-related neurons (see **Materials and Methods**). Each dot represents a task-related (TR; *black*) or non-task-related (NTR; *gray*) neuron. *Right*, Peri-event time histograms (PETHs) of spike activity in representative TR and NTR neurons (a–c). ***C***, Summary of the z-scored spike rate changes for the four clusters (Clu.1–4) of all TR (RS and FS) neurons, aligned with Go cue and reward of true consummation trials, and Go cue and no reward of false consummation trials (from left to right). Neurons were sorted based on the spike peak in the second alignment (true, reward). ***D***, Spike rate changes and task event relevance for the four neuron clusters. *Each top*, Z-scored spike rate change (mean and s.e.m.), aligned with the Go cue (*left*) and reward (*right*). *Each middle*, Onset-to-offset time-course of the spike activity, temporally normalized (*) by the interval between the Go cue and release (*left*) and the Correct cue and reward (*right*). *Each bottom*, Phase distribution in the PS analysis for the offset (Clu.1,2) and onset (Clu.3,4).

The spike activity of Clu.1 neurons characteristically decreased after the Go cue (or pedal release) (**Fig. 5*A***; Kruskal–Wallis test *p* = 2.55 × 10^−5^, *χ^2^* = 24.0, *η²* = 7.43 × 10^−3^; post hoc Dunn test, True-pre vs. True-post *p* = 7.79 × 10^−6^, etc.) and reward delivery (*p* = 5.45 × 10^−34^, *χ^2^* = 158, *η²* = 0.0524; True-pre vs. -post *p* = 2.73 × 10^−14^) in true consummation trials. The activity of Clu.2 decreased only after the reward delivery (*p* =1.38 × 10^−99^, *χ^2^* = 461, *η²* = 0.103; True-pre vs. -post *p* < 1.00 × 10^−300^). Notably, these activity changes in the Clu.1 and Clu.2 neurons, almost half of all task-related neurons, resembled the time-course of SWR occurrence. Conversely, Clu.3 neurons showed rapid spike increases in response to the Go cue/release (*p* = 4.74 × 10^−10^, *χ^2^* = 46.4, *η²* = 0.0125; True-pre vs. -post *p* = 2.12 × 10^−7^) and reward delivery (*p* = 8.40 × 10^−31^, *χ^2^* = 143, *η²* = 0.0394; True-pre vs. -post *p* < 1.00 × 10^−300^), while the Clu.4 neurons increased the activity much more slowly after the reward delivery (*p* = 1.02 × 10^−11^, *χ^2^* = 54.2, *η²* = 0.0190; True-pre vs. -post *p* = 1.10 × 10^−9^). Population analysis for each neuron cluster (**Fig. 4*D***) revealed that Clu.1 and Clu.3 neurons changed their spike activity between the Go cue and release. In contrast, Clu.1–3 neurons at once responded to reward delivery rather than to the Correct cue. The temporal profiles of task-related neurons were in good harmony with those of the SWRs (**Fig. 3*D***).

In addition, we evaluated how the activity patterns of these neuron clusters were affected by reward expectation and acquisition in the FTC task, using RMI (pre Go cue) and RMI (post reward), respectively (see **Materials and Methods**). The RMI (pre Go cue) of Clu.1 and Clu.2 neurons was biased positively (**Fig. 5*B***), meaning their activity was enhanced by reward expectation in the pedal hold period, as expected from the spike rate differences (**Fig. 5*A***; Go cue, False-pre vs. True-pre *p* = 3.00 × 10^−3^ for Clu.1 and *p* = 1.20 × 10^−5^ for Clu.2). In contrast, the RMI (post reward) of Clu.1 and Clu.2 neurons was remarkably biased towards the −1 end (**Fig. 5*B***), due to spike rate decrease after reward acquisition (**Fig. 5*A***; Reward, False-post vs. True-post *p* < 1.00 × 10^−300^ for Clu.1 and *p* < 1.00 × 10^−300^ for Clu.2). It is worth noting that the distribution of the SWRs overlapped with the protrusive part of the Clu.1 and Clu.2 neurons.

### Synchrony between SWRs and neuronal activities

Finally, we examined how hippocampal neurons with different functions behave during SWRs. As known well, many RS and FS neurons displayed spike increase in synchrony with SWRs (**Fig. 6*A***). Overall, the percentage of RS neurons showing SWR-coincided spike increases was significantly higher than that of FS neurons (**Fig. 6*B***; 2×4 *χ^2^* test *p* = 2.37 × 10^−75^, *χ^2^* = 349, *V* = 0.237). The percentage of task-related neurons with SWR-coincided spike increases was higher than that of non-task-related neurons (**Fig.6*C***; 2×4 *χ^2^* test *p* = 3.49 × 10^−83^, *χ^2^* = 385, *V* = 0.249). The peak distribution of significant spike increases was centered in the peak time of SWRs, while the trough distribution of spike decreases, seen only rarely, was delayed from the peak of the SWRs (**Fig. 6*D***; spike increase −5.0 ± 25.0 ms; decrease 23.0 ± 77.0 ms). In each cycle of ripples, the FS neurons discharged following the RS neurons (**Fig. 6*D***; RS 72.3 ± 44.4°; FS 119 ± 59.7°, Cramér-von Mises test *p* < 1.00 × 10^−300^, *P* = 70.8, *η²* = 0.0387). These results agree well with those of previous reports (Ylinen et al., 1995; Csicsvari et al., 1999; Klausberger et al., 2003), confirming that our SWR and spike data were sufficiently reliable for the analysis.

**Figure 5.**
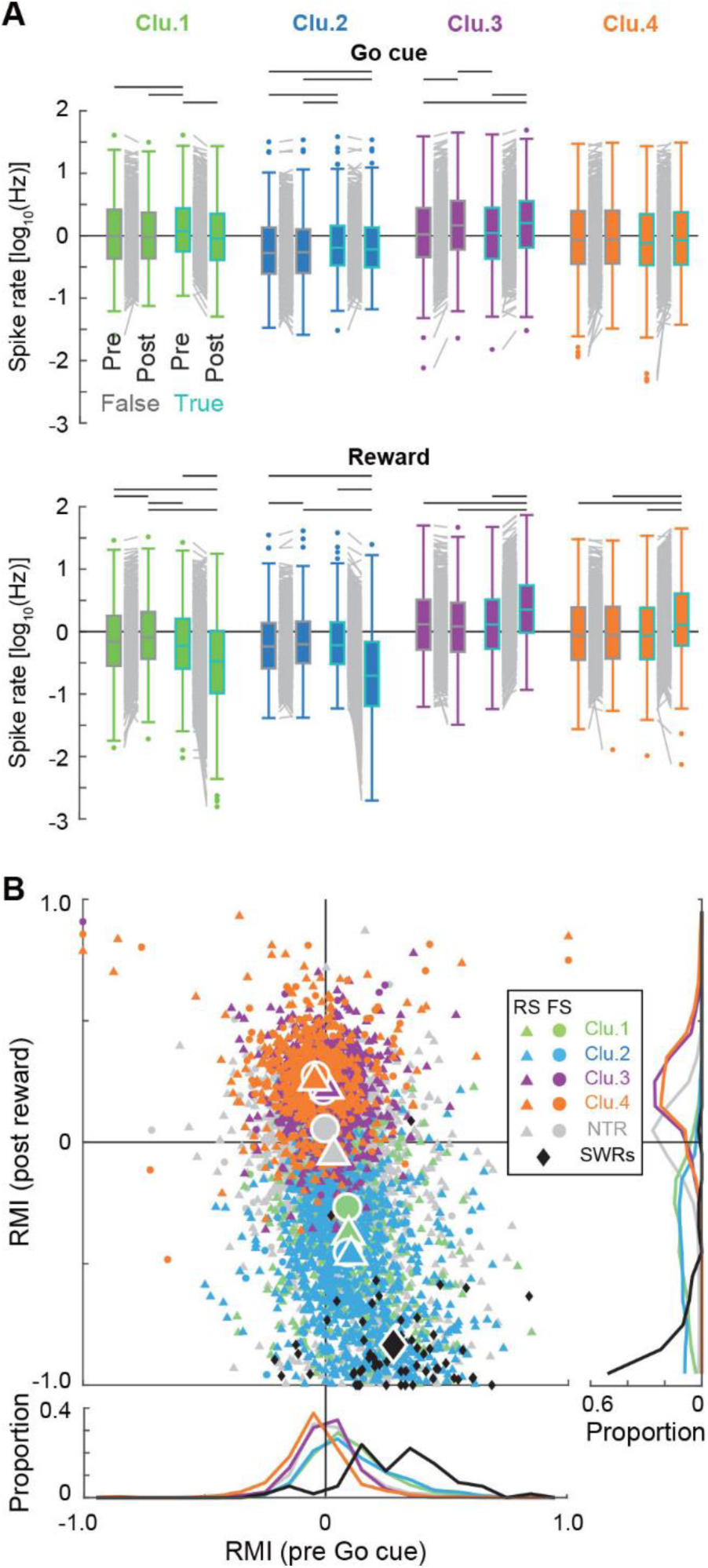
Modulation of neural activity by reward expectation and acquisition. ***A***, Comparison of the spike rates (logarithmic scale) between the pre- and post-event time windows (*upper*, Go cue; *lower*, Reward; 1 s) and between false and true consummation trials in individual neuron clusters (1–4). The horizontal lines represent statistical significance. ***B***, The reward modulation indices (RMIs) in the pre Go cue (*abscissa*) and post reward (*ordinate*) time windows. Each symbol represents a single neuron (*triangle*, RS; *circle*, FS) colored according to its cluster (including the NTR in gray). The black diamond represents the SWRs in each session. Large symbols indicate their medians. The RMI denotes higher (towards +1) or lower (−1) activity depending on reward expectation (pre Go cue) and acquisition (post reward). Note that the SWRs (*black*) overlap with the protrusive clu.1 (*green*) and 2 (*blue*) neurons.

**Figure 6.**
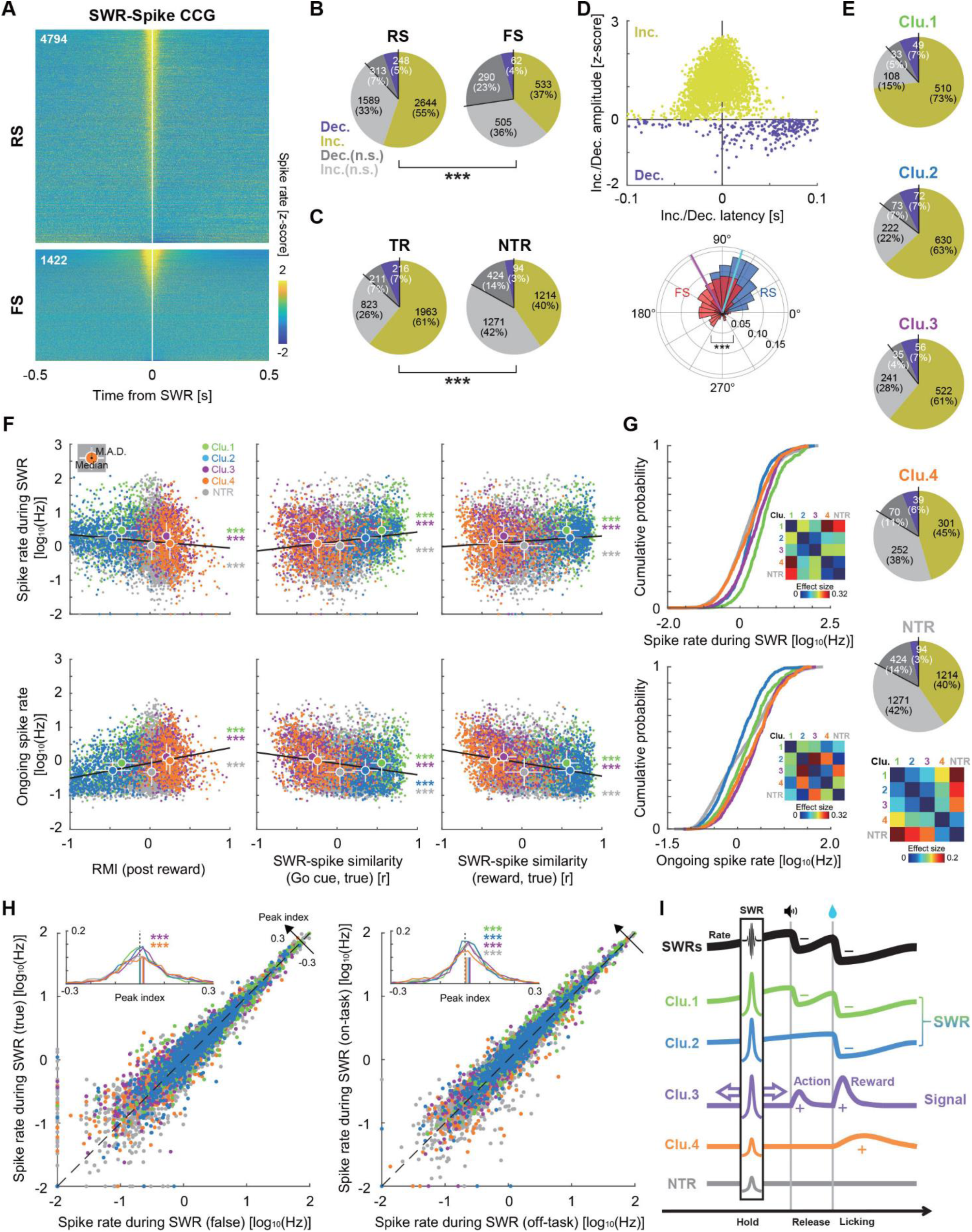
Spiking activity in synchrony with SWRs during FTC task performance. ***A***, Z-scored cross-correlogram (CCG) of spikes to the peak of SWRs (0 s) in individual neurons (all RS and FS neurons). ***B***, Proportion of spike increases (Inc.; significant in yellow, not significant in light gray) and decreases (Dec.; significant in dark purple, not significant in dark gray) during the SWRs in all RS and FS neurons. The RS neurons showed significantly more SWR synchrony than FS neurons did. Neuron counts and percentages are shown in each pie chart. ***C***, The proportion of spike increases and decreases during SWRs (with and without statistical significance) in all TR and NTR neurons. TR neurons showed more SWR synchrony than NTR neurons. ***D***, *Top*, Peak/trough latency of the spike increases/decreases from the SWR peak time (0 s). *Bottom*, Distribution of the most active phases of ripple oscillations in RS (*blue*) and FS (*red*) neurons. ***E***, Proportion of spike increases and decreases during SWRs in Clu.1–4 neurons and NTR neurons. Most Clu.1–3 neurons exhibited a spike increase synchronous with SWRs, with fewer in Clu.4 and NTR, as shown in the color matrix of statistical effect sizes for all pairs of clusters (*bottom*). ***F***, Plots of the spike rate (logarithmic scale) during SWRs (*upper*; ±50 ms from the SWR peak) and ongoing spike rate (*lower*) against the RMI in the post reward (*left*), similarity (correlation coefficient *r*) between SWR and spike rate changes around the Go cue (*middle*) and reward (*right*) along the course of true consummation trials. Each dot represents a single neuron colored by a cluster (Clu.1–4 and NTR) with a regression line for all dots (*black*). The large dots and bars indicate the median and median absolute values (M.A.D.). Colored asterisks indicate that the neuron cluster deviated significantly above or below the regression line (sign test). ***G***, Cumulative distributions of the spike rate during SWRs (*upper*) and the ongoing spike rate (*lower*) shown in ***F***. *Insets*, color of the effect size matrices for all cluster pairs. ***H***, *Left*, Plots of spike rates during SWRs between false and true consummation trials. The inset shows shifts towards higher peak index in the true consummation trials. *Right*, Plots of the spike rate during SWRs between off-task (outside task engagement) and on-task (during engagement) periods. The inset shows the shifts towards higher peak index during the on-task period. ***I***, Schema of hippocampal SWR and spike activity patterns. SWR and spike rate changes along an action-outcome trial (*traces*) and SWR occurrence with spike increases (*box*). Thick arrows represent possible replay and preplay of the Clu.3 signals. See our hypothesis in the **Discussion**.

**Figure 6*E*** summarizes the spike increases and decreases synchronized with the SWRs in the four clusters of task-related neurons and non-task-related neurons. The Clu.1 to Clu.3 neurons had more SWR-coincided spike increases compared with the Clu.4 and non-task-related neurons (5×2 *χ^2^* test *p* = 2.67 × 10^−102^, *χ^2^* = 514, *V* = 0.203). The actual spike rate during the SWRs was negatively correlated with the RMI (post reward), but positively correlated with the SWR-spike similarity (around Go cue and reward in true consummation trials; see **Materials and Methods**) overall for all the neurons [**Fig. 6*F***, *upper*; spike rate during SWR against RMI (post reward) *p* = 3.21 × 10^−18^, *r* = −0.110; against SWR-spike similarity (Go cue, true) *p* = 1.49 × 10^−39^, *r* = 0.166; against SWR-spike similarity (reward, true) *p* = 4.73 × 10^−17^, *r* = 0.106], suggesting functional correlates of the SWR synchrony and task-related activity pattern in hippocampal neurons. The ongoing spike rate also correlated with the RMI and SWR-spike similarities [**Fig. 6*F***, *lower*; ongoing spike rate against RMI (post reward) *p* = 1.01 × 10^−106^, *r* = 0.273; against SWR-spike similarity (Go cue, true) *p* = 3.53 × 10^−64^, *r* = −0.212; against SWR-spike similarity (reward, true) *p* = 1.23 × 10^−114^, *r* = −0.283]. Compared with the overall trends (regression lines), Clu.1 and Clu.3 neurons always showed higher spike rates, whereas non-task-related neurons showed lower spike rates (**Fig. 6*F***; sign test). Likewise, cumulative frequency curves showed that the Clu.1–3 neurons exhibited significantly stronger activation than the Clu.4 and non-task-related neurons (**Fig. 6*G***; *upper*; Kruskal–Wallis test *p* = 5.07 × 10^−79^, *χ^2^* = 371, *η²* = 0.0558; post hoc Dunn test, *p* < 0.05 among all cluster pairs except the Clu.4 and NTR combination), which supports the above observations (**Fig. 6*C*,*E*,*F***). On the other hand, the ongoing spike rate, reflecting baseline excitability, was highest in the Clu.3 and Clu.4 neurons, and lowest in the Clu.2 and non-task-related neurons (**Fig. 6*G***; *lower*; *p* = 3.52 × 10^−76^, *χ^2^* = 358, *η²* = 0.0539; *p* < 0.01 among all cluster pairs except the Clu.2 and NTR pair).

Furthermore, spike rates during the SWRs of the Clu.3 and Clu.4 neurons were slightly higher in true than false consummation trials (**Fig. 6*H***, *left*; Clu.3, one-sample Wilcoxon signed-rank test *p* = 6.84 × 10^−10^, *z* = 6.17, *r* = 0.211; Clu.4 *p* = 7.82 × 10^−5^, *z* = 3.95, *r* = 0.154), and those of Clu.1–3 neurons were also higher in the on-task than off-task period (**Fig. 6*H***, *right*; Clu.1 *p* = 6.19 × 10^−13^, *z* = 7.20, *r* = 0.272; Clu.2 *p* = 7.77 × 10^−11^, *z* = 6.51, *r* = 0.206; Clu.3 *p* = 1.15 × 10^−15^, *z* = 8.01, *r* = 0.274; NTR *p* = 1.51 × 10^−4^, *z* = 3.79, *r* = 0.069). Thus, the Clu.1–3 neurons displayed higher spike synchrony with the SWRs than the Clu.4 and non-task-related neurons, especially in the on-task period and true consummation trials.

## Discussion

In freely moving animals, the hippocampus drives theta oscillations that encode spatial information *online* during exploration, and SWRs that reproduce the information *offline* when they stop (Foster and Wilson, 2006; Diba and Buzsaki, 2007). At the time, the occurrence of SWRs is facilitated by the presence of rewards after the exploratory action (Singer and Frank, 2009; Ambrose et al., 2016; Sosa and Giocomo, 2021). Thus, it is now widely accepted that SWRs occur during consummatory behaviors, such as eating and drinking (Buzsáki 2015; Buzsáki and Tingley, 2023), in addition to simple immobility (O’Neill et al., 2006; Vöröslakos et al., 2023). However, in our immobile rats, SWRs disappeared during the consummatory licking action triggered by the presentation of reward water in the operant learning task (**Figs. 2,3**). SWR disappearance due to licking was similarly observed even in immobile rats without any reward learning (Fig. 1; cf., already referred to by Klee et al., 2021, supplemental data, for classical conditioning). In contrast, SWR occurrence increased with reward expectation during the preparatory pedal-holding period (**Figs. 2,3**). Further, one group of hippocampal neurons (Clu.3) responded to the Cue/release and reward, while others (Clu.1,2) showed similar characteristics to the time-course of SWR occurrence as described above (**Figs. 4,5**). These neurons discharged more synchronously with SWRs than with task-unrelated neurons during the on-task period (**Fig. 6**). Taken together, the established theory that SWRs occur during consummatory behaviors (as either consummation or consumption) does not seem applicable to immobile animals drinking under head fixation and body covering.

Consummatory behaviors other than eating and drinking, as defined psychologically (Craig, 1918; Woodworth, 1918), would also be informative in understanding the two exclusive modes of hippocampal oscillatory activity (i.e., theta oscillations *online* and SWRs *offline*). One such example is animal copulation. Both male ejaculation and female lordosis after ejaculation are considered consummatory behaviors that satisfy sexual desires. At this stage, theta oscillations, but not SWRs, are observed in the hippocampi of male (Kurtz and Adler, 1973) and female rats (Kurtz, 1975). Another example is voluntary wheel running in rodents, which can be a consummatory behavior in itself as a natural reward that satisfies the desire (presumably analogous to “exercise” for humans) (Brené et al., 2007; Basso and Morrell, 2015; Buhler et al., 2023). During this behavior, theta oscillations, but not SWRs, are observed in the hippocampus (Buhler et al., 2023). Thus, according to the classical psychological definition, both modes of hippocampal oscillatory activity may participate in consummatory behaviors that satisfy desires (appetite, sex, sleep, etc.). In other words, the idea that hippocampal SWRs occur during consummatory behaviors is difficult to generalize to all types of consummatory behaviors.

Given these facts, it is worthwhile to consider why SWRs behave differently during the same consummatory licking action under freely moving and head-fixed conditions. We may presume that SWRs are prone to appear during no or relatively little movement, that is, at a pause before and after physical actions, independent of behavioral consummation. In freely moving conditions, running involves heavy whole-body movements, whereas stopping to eat or drink results in relatively less movement. Such “relative immobility” (or action pause) may bring about more SWRs in the hippocampus. Under head-fixed immobilization, the entire body is kept motionless; thus, theta oscillations are hardly driven in the hippocampus. Although the pedal hold action is part of preparatory behavior, it is relatively immobile without limb and tongue movements, thereby allowing SWR generation in the hippocampus. Then, the SWRs are temporarily diminished by pedal release that reduces immobility. Once a real reward is delivered, the SWRs disappear due to a further reduction in immobility by intentionally drinking the reward. In fact, we have recently observed that the SWR occurrence dropped when any part of the whole body (jaw, nose, whiskers, limbs, trunk, tail, etc.) moved physically in similarly immobilized rats (Rios et al., 2024). As such, relative immobility may better explain the difference in SWR occurrence between the freely moving and head-fixed conditions.

Upon closer inspection, the SWRs disappeared sharply at the time of reward delivery itself, rather than during anticipatory/intake licking or additional pedal release (**Fig. 3D,*E***). In addition, the Correct cue prior to the reward did not affect the occurrence of SWRs (**Fig. 3D**). Therefore, the intrinsic factor that breaks SWR occurrence is likely to be *internal* information on the *intended* action, rather than actual downstream muscular movements or upstream perceptual/motivational processes. We consequently speculate that animals may rehearse and review each action through hippocampal SWR activity (preplay, Davidson et al., 2009; Gupta et al., 2010; Dragoi and Tonegawa, 2011; replay, Foster and Wilson, 2006; Diba and Buzsaki, 2007) at action pauses before and after the intended action, as if they thought then moved, stopped then thought. Maybe it is a natural and ongoing property of ordinary brain operations, as recently reported about the correlation between SWRs and internal thought patterns in human daily life (Iwata et al. 2024). Therefore, it is possible that immobile animals may also produce functional SWRs for the rehearsal and review of their intended actions.

Herein, we propose a hypothesis to account for the neuronal mechanism of action-outcome association under our experimental conditions (**Fig. 6*I***). First, the hippocampal neurons that play a direct role in task performance are the Clu.3 neurons that convey phasic signals of the action and its outcome reward. Our previous study showed that such hippocampal neurons emerge spike activity related to action and reward in the progress of operant learning (Soma et al. 2023). Unlike Clu.3 neurons, Clu.1 and Clu.2 neurons may work together with SWR occurrence during the pedal hold period, depending on the degree of reward expectation. It is possible that Clu.1 and Clu.2 neurons participate in the framework of SWR activity by elevating their excitability. If so, once an SWR occurs through Clu.1 and Clu.2 neurons before the action, Clu.3 neurons may also become active for *offline* reviewing the previous trial or rehearsing the current trial. Perhaps the SWRs occurring after completion of reward acquisition would have a similar effect on these neurons. The Clu.3 neurons themselves might have grown from action-related neurons and reward-related neurons (Soma et al. 2023) as a result of their associative spiking driven by the SWRs in the course of this operant learning. Thus, our hypothesis explains the process of association between the action and outcome.

As shown in freely moving animals (Singer and Frank, 2009; Ambrose et al., 2016; Sosa and Giocomo, 2021), reward expectation enhances SWR occurrence as well as Clu.1 and Clu.2 neuronal activities during the pedal hold period in our immobile rats. This would be advantageous for effectively increasing opportunities to reinforce the association between actions and outcomes. Interestingly, we found that reward expectation enhances neuronal activity related to the Go cue and push-pull actions in the dopamine reward system of the basal ganglia (Rios et al. 2023). This would suggest that the dopamine reward system can affect Clu.3 neurons rather than the SWRs and Clu.1/Clu.2 neurons in the hippocampus. The basal ganglia may facilitate the optimization of the action based on its outcome by repetitively interacting with the SWR-involved episodic memory system in the hippocampus, which may also contribute to the action-outcome association (Corbit and Balleine, 2000).

Overall, this study provides evidence that physical action during whole-body immobility critically determines the ability to generate SWRs in the hippocampus. Currently, head fixation protocols are widely applied in many neuroscience experiments using rodents and primates because they have practical advantages not only for accurate behavioral assessment of limbs, tongue, and whiskers, but also for electrical and optical measurement and manipulation of neural activity at high spatial and temporal resolutions. If such useful head fixation could be combined with virtual reality, which mimics visual or auditory perception in a freely moving context, cognitive brain functions could be causally tested, even in unreal circumstances (e.g., Harvey et al. 2009). Furthermore, brain status during head and body immobility may be relevant, not only to experimental animal behaviors, but also to human daily life. Consistent with our observations, Iwata et al. (2024) have recently reported that SWRs decrease during meals in the hippocampi of human subjects. Then, questions familiar to our daily life arise, for example, whether the hippocampus would adopt different modes for water drinking while jogging from while sitting in an airplane seat, or whether the hippocampus performs cognitive and memory functions differently in bedridden patients from ambulatory patients. Mobility and immobility are the basic determinants of brain function. Our findings shed new light on the dichotomous mechanisms of hippocampal oscillatory activity in cognition and memory by interpreting the behavioral significance of actions and action pauses in distinct physical situations.

## Acknowledgments

This work was supported by Grants-in-Aid for Scientific Research (B) (JP19H03342 and JP23H02589 to Y.I.), for Scientific Research on Innovative Areas (JP20H05053 to Y.I.), for Transformative Research Areas (A) (JP21H05242 to Y.I.), for Challenging Research (Exploratory) (JP24K21999 to Y.I.), and for Early-Carrer Scientists (JP22K15222 to M.K.) from MEXT and JSPS; by Brain/MINDS (JP19dm0207089 to Y.I.) from AMED; by CREST (JPMJCR1751 to Y.I.) and SPRING (JPMJSP2120 to T.S.) from JST; by the Takeda Science Foundation (Y.I.); and by Center for Brain Integration Research, Institute of Science Tokyo. We wish to express our appreciation to all (former) members of the Isomura laboratory, particularly Drs. Hidenori Aizawa, Akiko Saiki-Ishikawa, Toshikazu Samura, as this study was inspired by their earlier preliminary observations (Samura T et al. *Neuroscience 2016*, Yokohama, Japan).

